# Therapeutic trial of anle138b in mouse models of genetic prion disease

**DOI:** 10.1101/2022.10.11.511739

**Authors:** Sonia M Vallabh, Dan Zou, Rose Pitstick, Jill O’Moore, Janet Peters, Derek Silvius, Jasna Kriz, Walker S Jackson, George A Carlson, Eric Vallabh Minikel, Deborah E Cabin

## Abstract

Phenotypic screening has yielded small molecule inhibitors of prion replication that are effective in vivo against certain prion strains but not others. Here we sought to test the small molecule anle138b in multiple mouse models of prion disease. In mice inoculated with the RML strain of prions, anle138b doubled survival and durably suppressed astrogliosis measured by live animal bioluminescence imaging. In knock-in mouse models of the D178N and E200K mutations that cause genetic prion disease, however, we were unable to identify a clear, quantifiable disease endpoint against which to measure therapeutic efficacy. Among untreated animals, the mutations did not impact overall survival, and bioluminescence remained low out to >20 months of age. Vacuolization and PrP deposition were observed in some brain regions in a subset of mutant animals, but appeared unable to carry the weight of a primary endpoint in a therapeutic study. We conclude that not all animal models of prion disease are suited to well-powered therapeutic efficacy studies, and care should be taken in choosing the models that will support drug development programs.

## Introduction

Prion disease is a fatal neurodegenerative disease caused by conformation conversion of cellular prion protein, PrP^C^, into a misfolded conformer known as scrapie prion protein or PrP^Sc1^. Therapeutics currently in development for prion disease aim to either lower PrP^C^ expression^2,3^ or bind PrP^C4^. Phenotypic screening in prion-infected mouse neuroblastoma cells has also yielded small molecules that inhibit PrP^Sc^ accumulation without targeting PrP^C5^. Of these small molecule leads, anle138b^6^ has recently cleared a Phase I clinical trial in healthy volunteers^7^ and is currently in clinical development for synucleinopathies (NCT04685265). Because anle138b was previously shown effective against the RML strain of prions in wild-type mice^6^, we sought to determine whether it would be effective in mouse models of genetic prion disease. The knock-in mouse lines ki-3F4-FFI^8^ and ki-3F4-CJD^9^ harbor mutations orthologous to D178N and E200K, two of the three most common^10^ causes of genetic prion disease in humans. Here, we sought to test anle138b administered orally in chow both to RML prion-infected mice and to these knock-in models.

## Methods

### Study design

All studies were conducted at McLaughlin Research Institute between June 2014 and June 2016 under IACUC approval MRI-GAC11. Compared to inoculated models of prion disease, we envisioned that a therapeutic trial in a genetic mouse model would serve two goals. First, we could evaluate efficacy across different prion strains or subtypes, which is important because some antiprion drug candidates have proven strain-specific^11^. Second, we could evaluate efficacy in a spontaneously sick model, which might offer a unique opportunity to modulate prion initiation in addition to prion replication, relevant because individuals at risk for genetic prion disease appear to be negative for pathological markers most of their lives^12,13^. We selected the ki-3F4-FFI and ki-3F4-CJD models because their mutations represent common causes of genetic prion disease in humans^14,15^ and they do not overexpress PrP^8,9^. In addition to the mouse orthologs of the D178N and E200K mutations, these knock-in alleles also contain the 3F4 epitope common to many mammals other than mice, which allows detection with the 3F4 antibody. To control for any impact this might have on disease outcomes, we used ki-3F4-WT mice, which also have this epitope but no mutation and do not develop spontaneous disease^8^, as a control group. As a disease endpoint we chose live animal bioluminescence imaging, wherein a Tg(Gfap-luc) transgene^16^ couples luciferase expression to astrocytosis. *Gfap* is among the earliest upregulated genes in prion-inoculated mice^17,18^, and live animal imaging provides a real-time readout of this pathology^19^; aged ki-3F4-FFI mice were likewise reported to develop strong GFAP immunoreactivity in cerebellum and thalamus^8^. Under the expectation that the onset of astrocytosis would occur at age 13±3 (mean±sd) months, a sample size of N=10 per group was calculated to provide 80% power (by 2-sided T test) to detect a 30% delay in onset; anle138b in RML prion-infected mice had been reported to delay disease by 30-97% depending upon the timepoint of treatment initiation^6^. Because mice would be subject to humane euthanasia, we also included survival as an endpoint. Finally, we included histopathology because spongiform change and PrP deposition are pathognomonic for prion disease. Animal care and histology scoring were performed unblinded.

### Animals

Prion inoculations utilized homozygous Tg(Gfap-luc) mice on an FVB/N background. Experimental knock-in mice were hemizygous Tg(Gfap-luc) and homozygous knock-in, resulting in a mixed background of FVB/N (from the Gfap-luc mice) as well as 129/Ola and C57BL/6 (on which the knock-in alleles were created). All cohorts were mixed sex.

### Procedures and monitoring

For the RML inoculation experiment, animals were inoculated intracerebrally with 10 µL of 1% brain homogenate from animals terminally sick with the RML strain^20^ of prions using a 26G Hamilton syringe. Knock-in animals were not inoculated. Live animal imaging was performed similar to the manner described previously^3^. Animals were injected intraperitoneally with 150 mg/kg D-luciferin (GoldBio) in a 15 mg/mL solution and imaged in an IVIS Lumina II In Vivo Imaging System (Perkin Elmer) at 10-20 minutes post-injection. Animals underwent daily monitoring plus welfare checks weekly, escalating to thrice weekly at 90 dpi for inoculated animals, with euthanasia upon: ≥20% weight loss from a 4-month baseline, inability to reach food or water, difficulty breathing, or dermatitis or fight wounds refractory to treatment. All animals, regardless of cause of death, were included in survival curves. Due to the retrospective nature of the analysis, there were a total of 6 animals for which records were incomplete: 2 (1 in the knock-in experiment and 1 in the RML inoculation experiment) for which the exact date of death was not recorded and was instead approximated based on last known observation of the animal, and an additional 4 (all in the knock-in experiment) for which records provided date of death but did not specify the cause or circumstances of death.

### Genotyping

Gfap-luc mice were genotyped using primer 9020 TCTCTAAGGAAGTCGGGGAAGC and primer 9021 CAGCGGGAGCCACCTGATAGCCTT, and running on 1% agarose with ethidium bromide for an expected product of 430 bp in the presence of the transgene. Knock-in mice were genotyped using primer 75 GAGCAGATGTGCGTCACCCAG, primer 77 GAGCTACAGGTGGATAACCCC, and primer 105 CAACATGAAGCATATGGCA. PCR PP1 using primers 75 and 77 gives a 204 bp product in wild-type mice and 218 bp in all three knock-in lines, resolvable with 3-4% TAE agarose, while heterozygous knock-ins yield a larger heteroduplex. PP38 PCR using primers 105 and 77 yields a product that digests with BbsI for ki-3F4-CJD yielding bands of 244 and 276 bp, digests with MfeI for ki-3F4-FFI yielding bands of 210 and 310, and digests with neither enzyme for ki-3F4-WT.

### Compound formulation

Anle138b (CAS No. 882697-00-9, Figure 1A) was provided by Dr. Armin Giese and was formulated at 2g/kg in mouse chow by SSNIFF (Soest, Germany). Knock-in animals were weaned onto facility chow and then switched to anle138b or matched placebo diets (SSNIFF) at age ∼2 months (range: 30-86 days). In the RML inoculation experiment, controls received facility chow while treated mice switched to anle138b diet immediately upon inoculation at 0 days post-inoculation (dpi).

**Figure 1.**
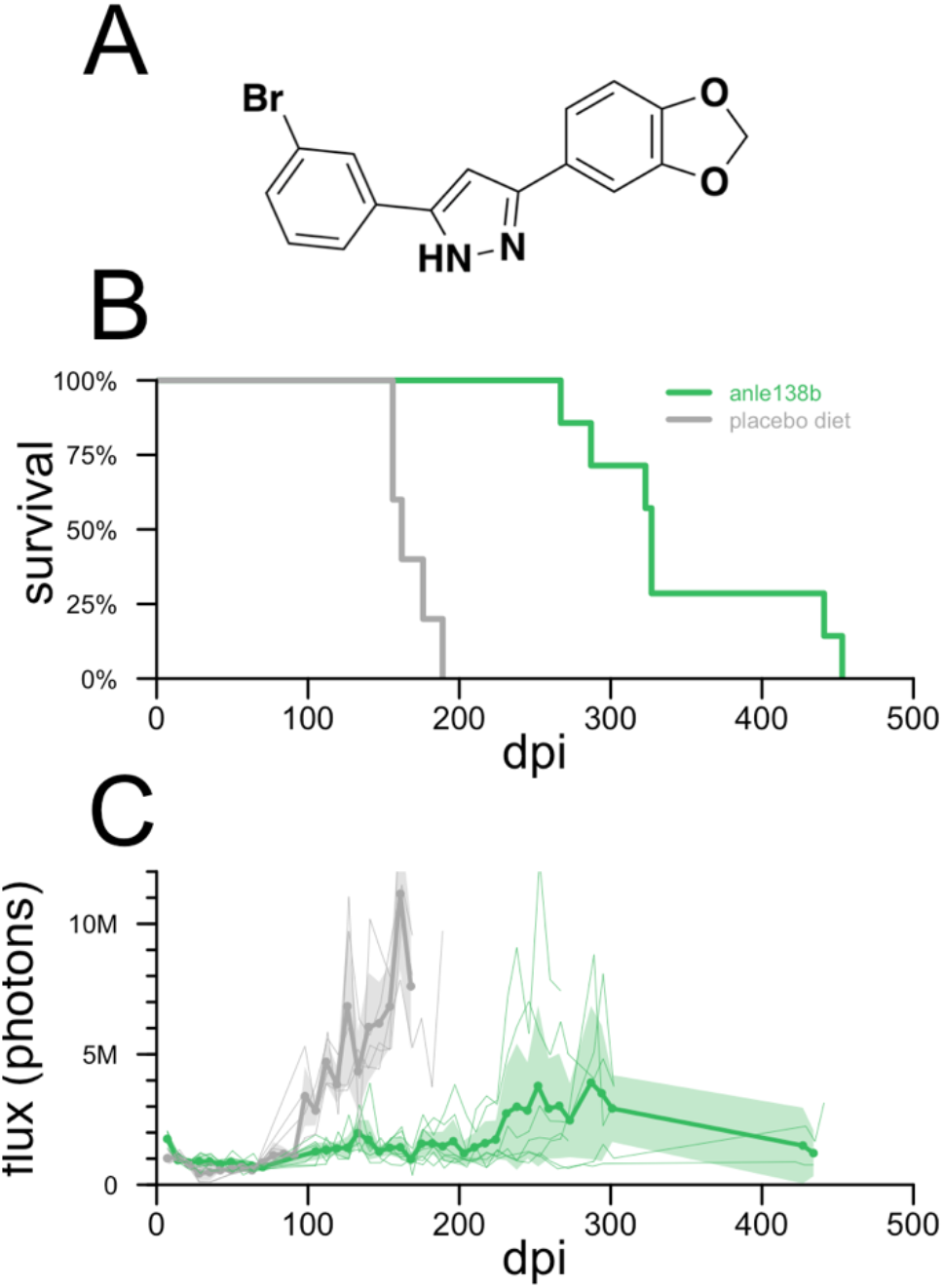
Efficacy of anle138b in RML prion-infected mice. **A)** Structure of anle138b. B) All-cause mortality for N=5 placebo diet and N=7 anle138b-treated Tg(Gfap-luc) mice with wild-type Prnp genes. C) Bioluminescence trajectories for the same animals. Thin lines indicate individual animal trajectories, thick lines and dots indicate means per week per treatment group, and shaded regions indicate 95% confidence intervals of the mean. Animals were imaged approximately weekly through ∼300 dpi; the two long-surviving treated animals were imaged again in two final sessions near end of life.

### Histology

Antigen retrieval was performed by boiling in 10mM sodium citrate pH 6.0 for 10 minutes. Primary antibodies were as follows. Iba1: rabbit monoclonal ab178846 (Abcam), 1:2000 3.5 minutes diaminobenzidine (DAB); secondary: biotinylated goat anti-rabbit IgG BA1000 (Vector Labs) 5:1000. PrP 3F4 primary: mouse monoclonal MAB1562 (Millipore) 1:1000 dilution, 5 minutes DAB; secondary: biotinylated goat anti-mouse IgG BA9200 (Vector Labs) 5:1000. We utilized VectaStain Elite ABC-HRP kit PK-6200 (Vector Labs), Permount mounting reagent SP15 (Fisher Scientific) and Mayer’s Hematoxylin S1275 (Sigma), and Eosin Y (Sigma HT110280).

### Statistics

All statistical analyses were conducted using custom scripts in R 4.2.0. All statistical tests were two-sided. Survival curves were compared using log-rank tests. Bioluminescence trajectories were compared using visualization of 95% confidence intervals; peak bioluminescence values were compared using Kolmogorov-Smirnov tests, which do not assume normality. Histology scores, because they are ordinal variables, were primarily compared using Wilcoxon tests, which do not assume normality and are applicable to rank data. Within each group of regions and endpoints being compared, false discovery rate (FDR) correction was used to control for multiple testing burden; FDR < 5% was considered significant. Because histology scores are ordinal data not well modeled by nested ANOVA, overall differences in histology scores across regions were additionally assessed using a cumulative link mixed model using the clmm2 function from the ordinal package in R^21^, with score (ordinal) modeled as a function of region, genotype, and, where applicable, treatment group (fixed effects) as well as individual animal (random effects).

### Source code and data availability

Source code and individual animal-level raw data sufficient to reproduce the figures and analyses herein are provided in a publicly available git repository at https://github.com/ericminikel/anle138b

## Results

In Tg(Gfap-luc) mice with wild-type *Prnp* genes inoculated with the RML strain of mouse prions, anle138b (Figure 1A) doubled survival (346±72 vs. 168±14 dpi, N=7 vs. 5, P = 0.00027, log-rank test, Figure 1B). The treatment also suppressed astrogliosis, delaying and in some animals preventing the increase in bioluminescence signal seen in control animals (Figure 1C); peak bioluminescence reached before death was 45% lower in treated animals (P = 0.015, Kolmogorov-Smirnov test).

Whereas RML-inoculated animals reach a terminal disease endpoint supportive of survival as a primary endpoint, the ki-3F4-FFI and ki-3F4-CJD knock-in models were reported to develop pathology in the absence of terminal illness^8,9^. We therefore prioritized histological and bioluminescence outcomes on the assumption that most animals would survive to end of study. Knock-in animals were followed out to 665-751 days of age before being harvested for histology. Among animals on placebo diet, overall survival assessed out to 719 did not differ among the three genotypes (P = 0.43, log-rank test, Figure 2A). In contrast to RML-inoculated mice (Figure 1C), bioluminescence for the three knock-in lines on placebo diet remained low and indistinguishable through 600 dpi (Figure 2B). At sacrifice, placebo-treated animals were scored for vacuolization and PrP puncta (3F4) staining across 9 brain regions (Figure 2C-D). After correction for multiple testing, ki-3F4-CJD animals exhibited 3F4 puncta scores significantly elevated above ki-3F4-WT controls in the lacunosum moleculare layer of the hippocampus as well as in olfactory granule cells (FDR < 5%, Wilcoxon test). No other differences were statistically significant at the group level, although light vacuolization could be observed in some ki-3F4-FFI and ki-3F4-CJD animals, particularly in the thalamus (Figure 3). 3F4 staining was overall faint in ki-3F4-FFI animals (Figure 3), consistent with underexpression of PrP and lack of clear deposition in this line^8^. Microglial activation assessed through Iba1 staining in select animals generally appeared to be at normal levels for aged animals and was not different between the knock-in lines (Figure 3). In a combined model accounting for individual animal and regional differences (see Methods), vacuolization was significantly enriched in ki-3F4-CJD animals (P=0.012 for ki-3F4-CJD, P = 0.060 for ki-3F4-FFI, each vs. ki-3F4-WT). Puncta likewise appeared enriched in ki-3F4-CJD animals (P value not defined because no puncta were observed in any ki-3F4-WT mice).

**Figure 2.**
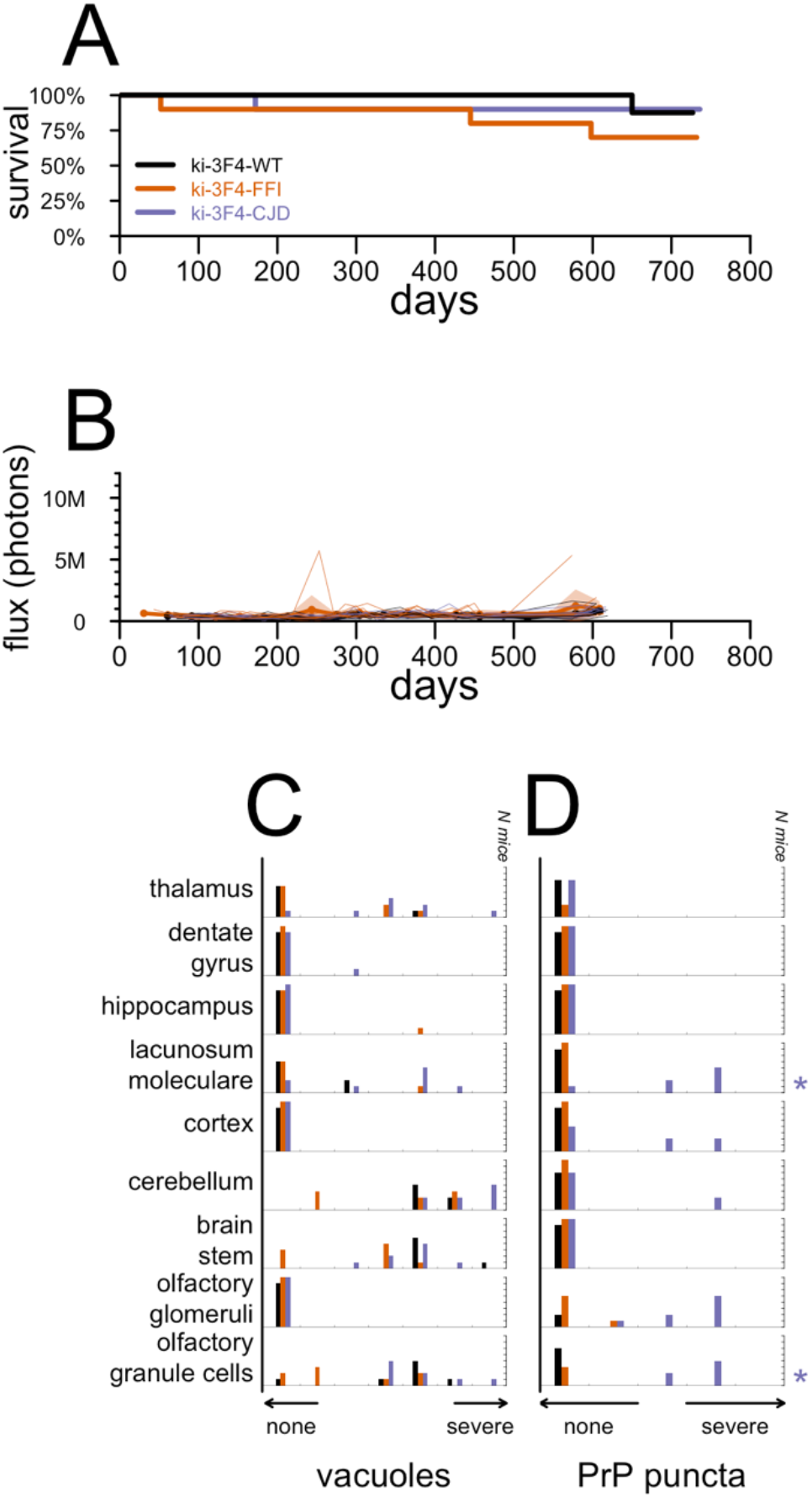
Candidate endpoints in placebo-treated knock-in mice. **A)** All-cause mortality for N=10 animals per group up through 719 dpi. **B)** Bioluminescence trajectories for the same animals. Animals were imaged approximately monthly, and occasionally twice monthly, through approximately 600 dpi. Thin lines indicate individual animal trajectories, thick lines and dots indicate means per week per treatment group, and shaded regions indicate 95% confidence intervals of the mean. For ease of comparison to RML prion-infected mice, the y axis scale is the same as Figure 1B. **C)** Histogram of vacuolization scores and **D)** histogram of 3F4 (PrP) puncta scores for N=5-8 animals per group subjected to histology analysis. Distributions for FFI and CJD mice were compared to WT mice for each region using Wilcoxon tests and multiple testing was corrected using false discovery rates. Symbols at right indicate corrected statistical significance: *FDR < 5%, **FDR < 1%, and are color-coded by genotype.

**Figure 3.**
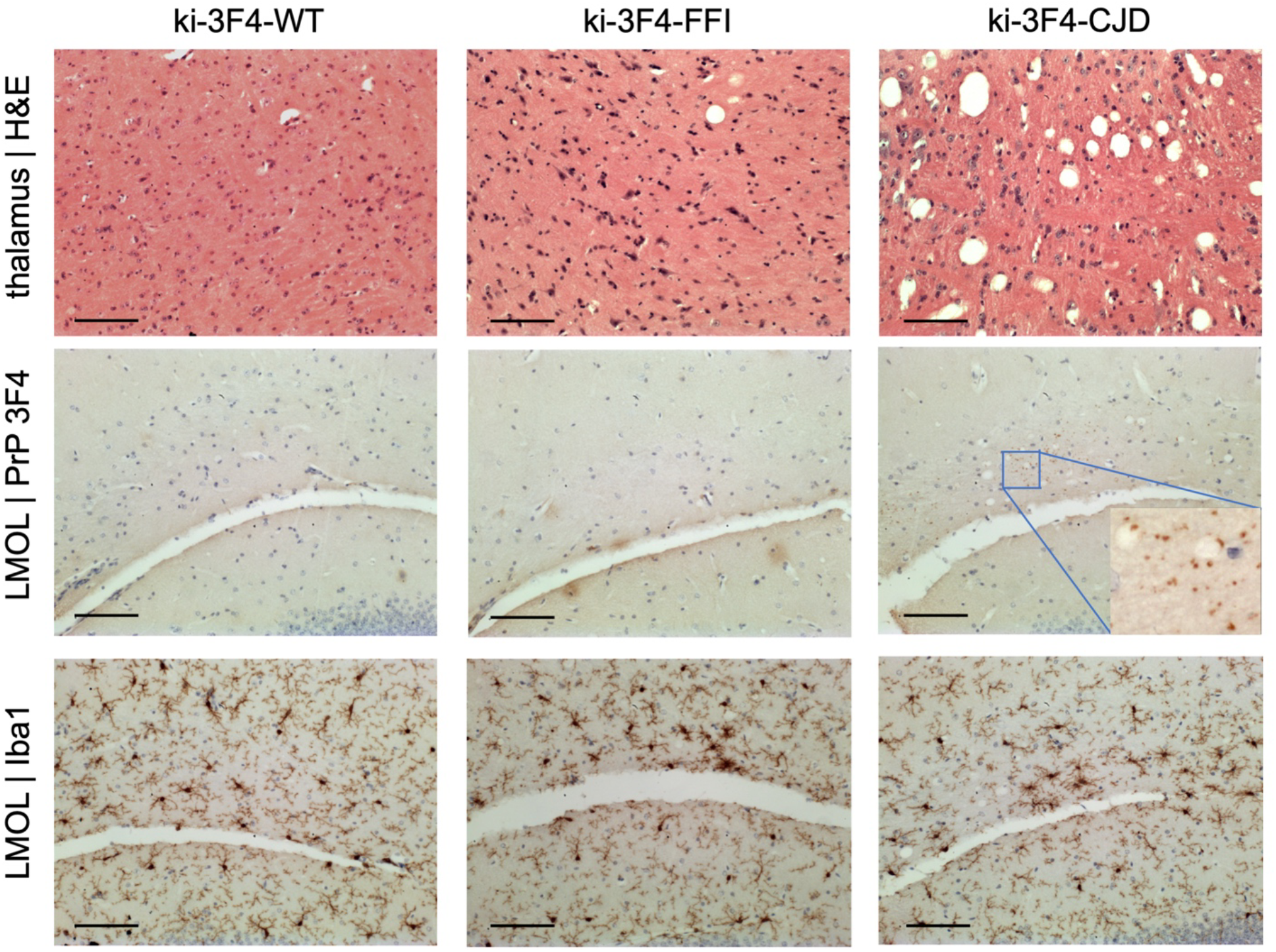
Example histological findings in placebo-treated knock-in mice. Rows show (top to bottom) vacuolization, 3F4 (PrP) immunoreactive puncta, and Iba1 (microglia); columns show one animal each of WT, FFI, and CJD placebo-treated groups. Scale bar: 10 µm.

Overall survival in knock-in animals treated with anle138b was indistinguishable from genotype-matched controls (P = 0.093 for ki-3F4-WT, Figure 4A; P = 0.23 for ki-3F4-FFI, Figure 4B; P = 0.15 for ki-3F4-CJD, Figure 4C; P=0.11 overall, log-rank test). Bioluminescence trajectories were similarly low for all groups, regardless of genotype and treatment group (Figure 4D, 4E, 4F). Histological analysis yielded no significant differences in vacuolization or PrP puncta when comparing treatment groups within genotypes (Figure 4G, 4H, 4I), and in a combined model (see Methods), neither outcome differed significantly by treatment group (P = 0.50 for vacuoles, P = 0.25 for puncta).

**Figure 4.**
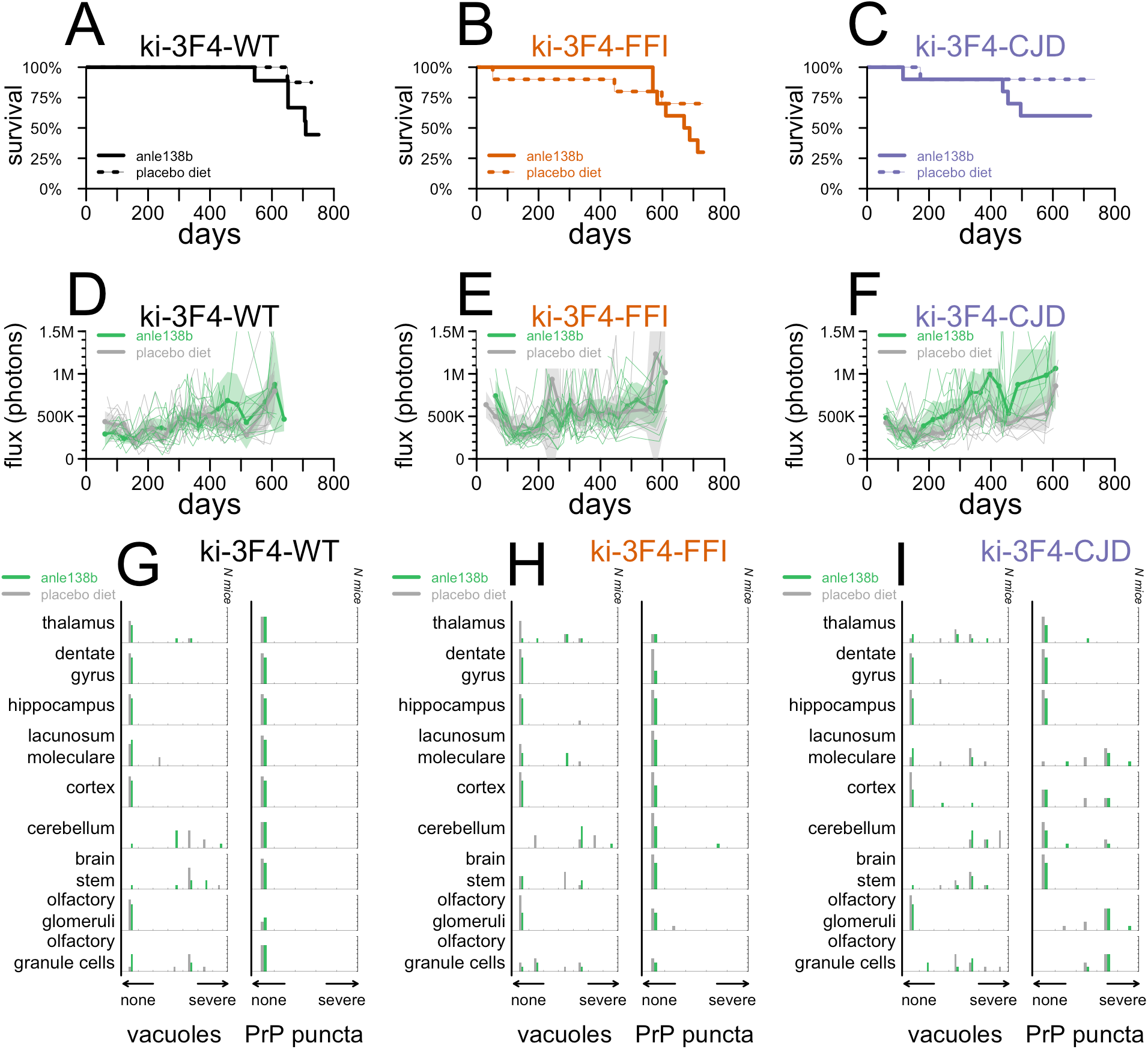
Effect of anle138b on endpoints in knock-in mice. **A-C)** All-cause mortality, **D-F)** bioluminescence trajectories, and **G-I)** histology endpoints in anle138b vs. placebo-treated animals of genotypes ki-3F4-WT **(A, D, G)**, ki-3F4-FFI **(B, C, H)**, and ki-3F4-CJD **(C, F, I)**.

## Discussion

Our pilot study in mice infected with the RML strain of prions replicated the survival benefit of anle138b treatment in this model^6^ and further demonstrated the treatment’s impact on a bioluminescence readout serving as a proxy for reactive astrogliosis. Interestingly, the response of prion disease-associated astrogliosis to therapy may depend upon the drug’s mechanism of action. In animals treated with a single dose of PrP-lowering ASO, astrogliosis remained low even as the drug washed out and animals eventually developed terminal prion disease^3^, and GFAP staining was reduced in treated animals that reached endpoint^2^. In contrast, the small-molecule PrP^Sc^ accumulation inhibitor cpd-B delayed the onset of astrogliosis, but treated animals ultimately reached bioluminescence levels equal to or even higher than untreated controls^22^. In the present study of anle138b, astrogliosis in treated animals remained low on average even as treated animals eventually succumbed to prion disease. This difference in averages is due in part, however, to the more variable time of onset among treated animals; some but not all treated animals did display a clear rise in bioluminescence before death. It will be interesting to determine whether and how distinct mechanisms of action of antiprion therapies might lead to differences in end-stage pathology.

PrP lowering therapy appears to be effective across prion strains^3^, whereas small molecules identified in through phenotypic screening have so far proven effective against some prion strains but not others^11,22–25^. Despite its efficacy in RML prion-infected mice, anle138b proved ineffective in humanized mice infected with sCJD MM1 prions^24^. In a mouse model of the A117V mutation, anle138b was reported to reduce plaque load without improving survival^26^. Here, we sought to test the efficacy of anle138b in two knock-in mouse models of genetic prion disease, corresponding to the E200K and D178N mutations. Unexpectedly, we were unable to evaluate the efficacy of anle138b because none of the selected endpoints provided a clear, quantitative measure of disease in these mice.

These knock-in mouse lines were previously reported to exhibit differences in body temperature regulation, burrowing, rotarod performance, and certain behavioral phenotypes extracted from video feeds by automated mouse behavioral analysis (AMBA), but a frank difference in all-cause mortality was not reported^8,9^, consistent with our findings here. A consistent increase in GFAP staining by 12-16 months was reported in these models^8,9^, inspiring our use of live animal imaging in Tg(Gfap-luc) mice as an endpoint here. However, subsequent studies did not identify a clear increase in Gfap mRNA levels in whole brain homogenates in these animals (Walker Jackson, personal communication), mirroring our results. Specific changes in mRNA translation have been observed in these knock-in lines^27^, distinct from those observed in RML-inoculated mice^28^, but these findings are more recent and were not leveraged here. Various histopathological abnormalities have also been identified in these knock-in mouse lines by 20 months (∼600 days)^8,9^, some of which we replicate here. Translating these histological changes into disease endpoints, however, is dependent upon staining and imaging conditions and on subjective scoring. It can be challenging to consistently translate these into quantitative endpoints. In addition, we cannot rule out that the penetrance or onset of pathologic features in the knock-in mice in this study may have been modified by their mixed genetic background, presence of the Gfap-luc allele, or myriad differences in facility or husbandry practices.

It has been proposed that antiprion therapeutics might enter clinical development in presymptomatic individuals at risk for genetic prion disease^29^. In this context efficacy studies in experimental models of genetic prion disease may seem especially appealing. A diversity of mouse models of genetic prion disease have been developed^30^, which exhibit many of the pathognomonic features of human prion disease, including spongiform change, PrP^Sc^ deposition, transmissibility, and/or fatality. The present study suggests that such models may in some cases yield invaluable insights into the basic science of prion disease without meeting additional requirements specific to the context of therapeutic studies. In particular, efficacy studies are served by disease endpoints that are quantitative, objective, easily measured, monitorable in real time, highly penetrant, and tightly distributed in severity or onset.

Given the difficulty of modeling most neurological diseases in animals, the diversity of models and even species available to prion research by virtue of the pan-mammalian nature of the disease is an unusual endowment. Though this panoply is united by the core prion disease process, the present study may serve as a cautionary reminder that, as we have argued elsewhere with regards to non-human primates^31^, not all prion models are ideally suited to all research purposes. Ultimately, multiple models will be needed to stitch together the complex continuum that bridges fundamental disease biology to therapeutic hypotheses and ultimately, drug development.

## Acknowledgments

This study was funded by 101 individual donors to an Experiment crowdfunding campaign (https://experiment.com/projects/can-anle138b-delay-the-onset-of-genetic-prion-disease), with additional support from Prion Alliance. The authors also acknowledge support from the National Institutes of Health (R01 NS125255 to SV and P01 NS041997 to GAC).

## Author contributions

Conceived and designed the study: SMV, GAC, WSJ, AG, EVM. Supervised the research: SMV, JK, GAC, EVM, DEC. Analyzed the data: SMV, EVM, DEC. Performed the experiments: DZ, RP, DS, JOM, DEC. Drafted the manuscript: SMV, EVM. Reviewed and approved the final manuscript: all authors.

## References

1. Prusiner SB. Prions. Proc Natl Acad Sci USA. 1998 Nov 10;95(23):13363–13383. PMCID: PMC33918

2. Raymond GJ, Zhao HT, Race B, Raymond LD, Williams K, Swayze EE, Graffam S, Le J, Caron T, Stathopoulos J, O’Keefe R, Lubke LL, Reidenbach AG, Kraus A, Schreiber SL, Mazur C, Cabin DE, Carroll JB, Minikel EV, Kordasiewicz H, Caughey B, Vallabh SM. Antisense oligonucleotides extend survival of prion-infected mice. JCI Insight. 2019 30;5. PMID: 31361599

3. Minikel EV, Zhao HT, L. J, O’Moore J, Pitstick R, Graffam S, Carlson GA, Kavanaugh MP, Kriz J, Kim JB, Ma J, Wille H, Aiken J, McKenzie D, Doh-Ura K, Beck M, O’Keefe R, Stathopoulos J, Caron T, Schreiber SL, Carroll JB, Kordasiewicz HB, Cabin DE, Vallabh SM. Prion protein lowering is a disease-modifying therapy across prion disease stages, strains and endpoints. Nucleic Acids Res. 2020 Aug 10; PMID: 32776089

4. Mead S, Khalili-Shirazi A, Potter C, Mok T, Nihat A, Hyare H, Canning S, Schmidt C, Campbell T, Darwent L, Muirhead N, Ebsworth N, Hextall P, Wakeling M, Linehan J, Libri V, Williams B, Jaunmuktane Z, Brandner S, Rudge P, Collinge J. Prion protein monoclonal antibody (PRN100) therapy for Creutzfeldt-Jakob disease: evaluation of a first-in-human treatment programme. Lancet Neurol. 2022 Apr;21(4):342–354. PMID: 35305340

5. Ghaemmaghami S, Russo M, Renslo AR. Successes and challenges in phenotype-based lead discovery for prion diseases. J Med Chem. 2014 Aug 28;57(16):6919–6929. PMCID: PMC4148153

6. Wagner J, Ryazanov S, Leonov A, Levin J, Shi S, Schmidt F, Prix C, Pan-Montojo F, Bertsch U, Mitteregger-Kretzschmar G, Geissen M, Eiden M, Leidel F, Hirschberger T, Deeg AA, Krauth JJ, Zinth W, Tavan P, Pilger J, Zweckstetter M, Frank T, Bähr M, Weishaupt JH, Uhr M, Urlaub H, Teichmann U, Samwer M, Bötzel K, Groschup M, Kretzschmar H, Griesinger C, Giese A. Anle138b: a novel oligomer modulator for disease-modifying therapy of neurodegenerative diseases such as prion and Parkinson’s disease. Acta Neuropathol. 2013 Jun;125(6):795–813. PMCID: PMC3661926

7. Levin J, Sing N, Melbourne S, Morgan A, Mariner C, Spillantini MG, Wegrzynowicz M, Dalley JW, Langer S, Ryazanov S, Leonov A, Griesinger C, Schmidt F, Weckbecker D, Prager K, Matthias T, Giese A. Safety, tolerability and pharmacokinetics of the oligomer modulator anle138b with exposure levels sufficient for therapeutic efficacy in a murine Parkinson model: A randomised, double-blind, placebo-controlled phase 1a trial. EBioMedicine. 2022 Jun;80:104021. PMCID: PMC9065877

8. Jackson WS, Borkowski AW, Faas H, Steele AD, King OD, Watson N, Jasanoff A, Lindquist S. Spontaneous generation of prion infectivity in fatal familial insomnia knockin mice. Neuron. 2009 Aug 27;63(4):438–450. PMCID: PMC2775465

9. Jackson WS, Borkowski AW, Watson NE, King OD, Faas H, Jasanoff A, Lindquist S. Profoundly different prion diseases in knock-in mice carrying single PrP codon substitutions associated with human diseases. Proc Natl Acad Sci USA. 2013 Sep 3;110(36):14759–14764. PMCID: PMC3767526

10. Goldman JS, Vallabh SM. Genetic counseling for prion disease: Updates and best practices. Genet Med. 2022 Jul 12;S1098-3600(22)00812–7. PMID: 35819418

11. Berry DB, Lu D, Geva M, Watts JC, Bhardwaj S, Oehler A, Renslo AR, DeArmond SJ, Prusiner SB, Giles K. Drug resistance confounding prion therapeutics. Proc Natl Acad Sci USA. 2013 Oct 29;110(44):E4160–4169. PMCID: PMC3816483

12. Vallabh SM, Minikel EV, Williams VJ, Carlyle BC, McManus AJ, Wennick CD, Bolling A, Trombetta BA, Urick D, Nobuhara CK, Gerber J, Duddy H, Lachmann I, Stehmann C, Collins SJ, Blennow K, Zetterberg H, Arnold SE. Cerebrospinal fluid and plasma biomarkers in individuals at risk for genetic prion disease. BMC Med. 2020 Jun 18;18(1):140. PMCID: PMC7302371

13. Thompson AGB, Anastasiadis P, Druyeh R, Whitworth I, Nayak A, Nihat A, Mok TH, Rudge P, Wadsworth JDF, Rohrer J, Schott JM, Heslegrave A, Zetterberg H, Collinge J, Jackson GS, Mead S. Evaluation of plasma tau and neurofilament light chain biomarkers in a 12-year clinical cohort of human prion diseases. Mol Psychiatry. 2021 Mar 5; PMID: 33674752

14. Minikel EV, Vallabh SM, Lek M, Estrada K, Samocha KE, Sathirapongsasuti JF, McLean CY, Tung JY, Yu LPC, Gambetti P, Blevins J, Zhang S, Cohen Y, Chen W, Yamada M, Hamaguchi T, Sanjo N, Mizusawa H, Nakamura Y, Kitamoto T, Collins SJ, Boyd A, Will RG, Knight R, Ponto C, Zerr I, Kraus TFJ, Eigenbrod S, Giese A, Calero M, de Pedro-Cuesta J, Haïk S, Laplanche JL, Bouaziz-Amar E, Brandel JP, Capellari S, Parchi P, Poleggi A, Ladogana A, O’Donnell-Luria AH, Karczewski KJ, Marshall JL, Boehnke M, Laakso M, Mohlke KL, Kähler A, Chambert K, McCarroll S, Sullivan PF, Hultman CM, Purcell SM, Sklar P, van der Lee SJ, Rozemuller A, Jansen C, Hofman A, Kraaij R, van Rooij JGJ, Ikram MA, Uitterlinden AG, van Duijn CM, Exome Aggregation Consortium (ExAC), Daly MJ, MacArthur DG. Quantifying prion disease penetrance using large population control cohorts. Sci Transl Med. 2016 Jan 20;8(322):322ra9. PMCID: PMC4774245

15. Minikel EV, Vallabh SM, Orseth MC, Brandel JP, Haïk S, Laplanche JL, Zerr I, Parchi P, Capellari S, Safar J, Kenny J, Fong JC, Takada LT, Ponto C, Hermann P, Knipper T, Stehmann C, Kitamoto T, Ae R, Hamaguchi T, Sanjo N, Tsukamoto T, Mizusawa H, Collins SJ, Chiesa R, Roiter I, de Pedro-Cuesta J, Calero M, Geschwind MD, Yamada M, Nakamura Y, Mead S. Age at onset in genetic prion disease and the design of preventive clinical trials. Neurology. 2019 Jun 6; PMID: 31171647

16. Zhu L, Ramboz S, Hewitt D, Boring L, Grass DS, Purchio AF. Non-invasive imaging of GFAP expression after neuronal damage in mice. Neurosci Lett. 2004 Sep 2;367(2):210–212. PMID: 15331155

17. Hwang D, Lee IY, Yoo H, Gehlenborg N, Cho JH, Petritis B, Baxter D, Pitstick R, Young R, Spicer D, Price ND, Hohmann JG, Dearmond SJ, Carlson GA, Hood LE. A systems approach to prion disease. Mol Syst Biol. 2009;5:252. PMCID: PMC2671916

18. Sorce S, Nuvolone M, Russo G, Chincisan A, Heinzer D, Avar M, Pfammatter M, Schwarz P, Delic M, Müller M, Hornemann S, Sanoudou D, Scheckel C, Aguzzi A. Genome-wide transcriptomics identifies an early preclinical signature of prion infection. PLoS Pathog. 2020 Jun 29;16(6):e1008653. PMID: 32598380

19. Tamgüney G, Francis KP, Giles K, Lemus A, DeArmond SJ, Prusiner SB. Measuring prions by bioluminescence imaging. Proc Natl Acad Sci USA. 2009 Sep 1;106(35):15002–15006. PMCID: PMC2736416

20. Chandler RL. Encephalopathy in mice produced by inoculation with scrapie brain material. Lancet. 1961 Jun 24;1(7191):1378–1379. PMID: 13692303

21. Christensen RHB. Cumulative link models for ordinal regression with the R package ordinal. [cited 2022 Sep 5]; Available from: http://cran.uni-muenster.de/web/packages/ordinal/vignettes/clm_article.pdf

22. Lu D, Giles K, Li Z, Rao S, Dolghih E, Gever JR, Geva M, Elepano ML, Oehler A, Bryant C, Renslo AR, Jacobson MP, Dearmond SJ, Silber BM, Prusiner SB. Biaryl amides and hydrazones as therapeutics for prion disease in transgenic mice. J Pharmacol Exp Ther. 2013 Nov;347(2):325–338. PMCID: PMC3807058

23. Berry D, Giles K, Oehler A, Bhardwaj S, DeArmond SJ, Prusiner SB. Use of a 2-aminothiazole to Treat Chronic Wasting Disease in Transgenic Mice. J Infect Dis. 2015 Jul 15;212 Suppl 1:S17–25. PMCID: PMC4551108

24. Giles K, Berry DB, Condello C, Hawley RC, Gallardo-Godoy A, Bryant C, Oehler A, Elepano M, Bhardwaj S, Patel S, Silber BM, Guan S, DeArmond SJ, Renslo AR, Prusiner SB. Different 2-Aminothiazole Therapeutics Produce Distinct Patterns of Scrapie Prion Neuropathology in Mouse Brains. J Pharmacol Exp Ther. 2015 Oct;355(1):2–12. PMID: 26224882

25. Giles K, Berry DB, Condello C, Dugger BN, Li Z, Oehler A, Bhardwaj S, Elepano M, Guan S, Silber BM, Olson SH, Prusiner SB. Optimization of Aryl Amides that Extend Survival in Prion-Infected Mice. J Pharmacol Exp Ther. 2016 Sep;358(3):537–547. PMCID: PMC4998675

26. Qin K, Zhao L, Solanki A, Busch C, Mastrianni J. Anle138b prevents PrP plaque accumulation in Tg(PrP-A116V) mice but does not mitigate clinical disease. J Gen Virol. 2019 Jun;100(6):1027–1037. PMID: 31045489

27. Bauer S, Dittrich L, Kaczmarczyk L, Schleif M, Benfeitas R, Jackson WS. Translatome profiling in fatal familial insomnia implicates TOR signaling in somatostatin neurons. Life Sci Alliance. 2022 Nov;5(11):e202201530. PMID: 36192034

28. Kaczmarczyk L, Schleif M, Dittrich L, Williams RH, Koderman M, Bansal V, Rajput A, Schulte T, Jonson M, Krost C, Testaquadra FJ, Bonn S, Jackson WS. Distinct translatome changes in specific neural populations precede electroencephalographic changes in prion-infected mice. PLoS Pathog. 2022 Aug;18(8):e1010747. PMCID: PMC9401167

29. Vallabh SM, Minikel EV, Schreiber SL, Lander ES. Towards a treatment for genetic prion disease: trials and biomarkers. Lancet Neurol. 2020 Apr;19(4):361–368. PMID: 32199098

30. Watts JC, Prusiner SB. Mouse models for studying the formation and propagation of prions. J Biol Chem. 2014 Jul 18;289(29):19841–19849. PMCID: PMC4106304

31. Mortberg MA, Minikel EV, Vallabh SM. Analysis of non-human primate models for evaluating prion disease therapeutic efficacy. PLoS Pathog. 2022 Aug 22;18(8):e1010728. PMID: 35994510

